# Uncultured marine cyanophages encode for active NblA, phycobilisome proteolysis adaptor protein

**DOI:** 10.1101/494369

**Authors:** Omer Nadel, Andrey Rozenberg, José Flores-Uribe, Shirley Larom, Rakefet Schwarz, Oded Béjà

## Abstract

Phycobilisomes (PBS) are large water-soluble membrane-associated complexes in cyanobacteria and some chloroplasts that serve as a light-harvesting antennas for the photosynthetic apparatus. When short of nitrogen or sulfur, cyanobacteria readily degrade their phycobilisomes allowing the cell to replenish the vanishing nutrients. The key regulator in the degradation process is NblA, a small protein (~6 kDa) which recruits proteases to the PBS. It was discovered previously that not only do cyanobacteria possess *nblA* genes but also that they are encoded by genomes of some freshwater cyanophages. A recent study, using assemblies from oceanic metagenomes, revealed genomes of a novel uncultured marine cyanophage lineage which contain genes coding for the PBS degradation protein. Here, we examine the functionality of *nblA*-like genes from these marine cyanophages by testing them in a freshwater model cyanobacterial *nblA* knockout. One of the viral NblA variants could complement the non-bleaching phenotype and restore PBS degradation. Our findings reveal a functional NblA from a novel marine cyanophage lineage. Furthermore, we shed new light on the distribution of *nblA* genes in cyanobacteria and cyanophages.

**Originality-Significance Statement:** This is the first study to examine the distribution and function of *nblA* genes of marine cyanophage origin. We describe as well the distribution of *nblA*-like genes in marine cyanobacteria using bioinformatic methods.

## Introduction

Phycobilisomes (PBS) are large light-harvesting protein supercomplexes (3-7 MDa) that are widely distributed in cyanobacteria, glaucocystophytes, and rhodophytes (Watanabe and Ikeuchi 2013; Adir 2005). These PBS antennas form water soluble protein-pigment complexes which are anchored to photosynthetic membranes. A typical PBS comprises several peripheral rods that project radially from the central core subcomplex. The central core consists of disks of allophycocyanin (AP) trimers and the peripheral rod subcomplexes consist of disks of phycobiliproteins [e.g. phycocyanin (PC), phycoerythrocyanin (PEC), or phycoerythrin (PE)], depending on the particular organism and the environmental conditions (Adir 2005).

It has been discovered that certain environmental conditions substantially affect PBS composition and the amount of cellular phycobiliproteins. For instance, the PBS can be adapted to changes in the ambient light by modifying the PBS pigment composition, a process called complementary chromatic acclimation (Gutu and Kehoe 2012; Montgomery 2017). Furthermore, nutrient limitation might lead to a complete degradation of the PBS (Collier and Grossman 1994). This is observed when cyanobacteria face nitrogen or sulfur starvation and results in cell bleaching. Currently, two biological functions are attributed to PBS degradation. First, dismantling of the PBS releases nutrients in the form of amino acids; second, reduced cellular metabolism under starvation might lead to over-excitation and therefore, degradation and uncoupling of the PBS serve as a mechanism to reduce electron transfer rate under such conditions. Multiple studies of PBS degradation have provided insight into the mechanism underlying this process (Forchhammer and Schwarz 2018).

Although the knowledge of the molecular mechanisms of PBS disassembly is very fragmentary, the key proteins involved in the process have been already known for two decades. In a genetic screen, mutants of *Synechococcus elongatus* were identified that were unable to degrade their PBS under nitrogen starvation (Collier and Grossman 1994). These non-bleaching mutants (*nbl*) retained their greenish color during nitrogen stress while the wild type displayed the typical yellowish bleached phenotype. Among the identified genes were *nblA* (Collier and Grossman 1994) and *nblB* (Dolganov and Grossman 1999). NblA is a small protein of ~60 aa with homologs found in many cyanobacteria, as well as in red algae (Nakamura et al. 2013; Stoebe et al. 1998). Biochemical assays, supported by the solved crystal structure, revealed that the functional form of NblA is a dimer (Bienert et al. 2006; Dines et al. 2008). NblA binds to the α-subunits of PC and PEC (Karradt et al. 2008) and the ß-subunit of PC (Nguyen et al. 2017), and in addition, it interacts with ClpC to recruit the Clp protease and induce proteolytic degradation of pigment proteins. Accordingly, it was suggested that NblA serves as a proteolysis adaptor, namely it interacts with the pigment and designates it to degradation (Dines et al. 2008; Karradt et al. 2008). Additionally, based on the close association of NblA with PBS complexes while attached to the photosynthetic membranes, NblA was also assigned a role in PBS disassembly (Sendersky et al. 2014; Sendersky et al. 2015). Although the cyanobacteria-derived chloroplasts of red algae also undergo bleaching under nitrogen starvation, they do not seem to recruit the plastid homologe of *nblA, ycf18*, for this function (Kawakami et al. 2009), although no experiments were performed on the other *nblA* homologe encoded in their nuclear genomes (Nakamura et al. 2013).

Over the last decade it became clear that many genes involved in photosynthesis are encoded in the genomes of viruses that infect cyanobacteria (cyanophages) and corresponding sequences were also identified in metagenomic assemblies originating from phages [see (Puxty et al. 2015) for review]. These auxiliary metabolic genes (AMGs) (Breitbart et al. 2007) seem to increase phage fitness during infection by supporting the growing metabolic demand needed to produce phage progeny (Lindell et al. 2005; Bragg and Chisholm 2008; Hellweger 2009). Analogously, genes encoding proteins required for bilin biosynthesis are also frequently found in cyanophages (Puxty et al. 2015). Moreover, it appears that cyanophage genomes might carry genes for PBS degradation, and *nblA*-like genes were described in genomes of some cyanophages infecting freshwater cyanobacteria (Gao et al. 2012; Ou et al. 2015). At the same time, *nblA*-like genes were not reported from marine cyanophages till now. This does not come as a surprise as their hosts are reported to lack *nblA* genes as well (Palenik et al. 2003; Karradt et al. 2008). Yet, this is contradicted by the fact that a number of *nblA* genes are actually annotated in marine cyanobacteria using automated annotation pipelines (e.g. http://www.ebi.ac.uk/interpro/entry/IPR007574/), from which one might hypothesize that marine cyanophages might potentially have them as well.

In the current paper, we report on the discovery of putative *nblA* genes from marine cyanophages with a focus on two genes found in a cyanophage genome obtained from metagenomic data. By expressing cyanophage-derived *nblA* genes in a freshwater model cyanobacterium, we show that one of the two proteins is able to restore the wild-type phenotype in such an assay. In addition, a systematic inventorization of *nblA* genes in annotated genomes reveals the existence of other marine cyanophages with *nblAs* and indicates that most PBS-containing cyanobacteria contain *nblA*-like genes in their genomes. We also highlight the fact that multiplicity of *nblA* genes per genome, while rather exceptional for cyanophages, is in reality a widespread phenomenon in cyanobacteria, and discuss the implications of these observations in the context of NblA complex structure.

## Results and discussion

Using metagenomic assembled genomes (MAGs) from the Red Sea (Philosof et al. 2017) and the *Tara* Oceans metagenomes (Brum et al. 2015; Sunagawa et al. 2015), we recently identified a novel uncultured oceanic cyanophage lineage (Flores-Uribe et al. 2018). Genomes of cyanophages belonging to this lineage contain common cyanophage genes, as well as genes coding for integrases, split DNA polymerases and NblA-like proteins (Flores-Uribe et al. 2018). Some of these phages appear to contain two different *nblA*-like genes (Figure 1a).

**Figure 1.**
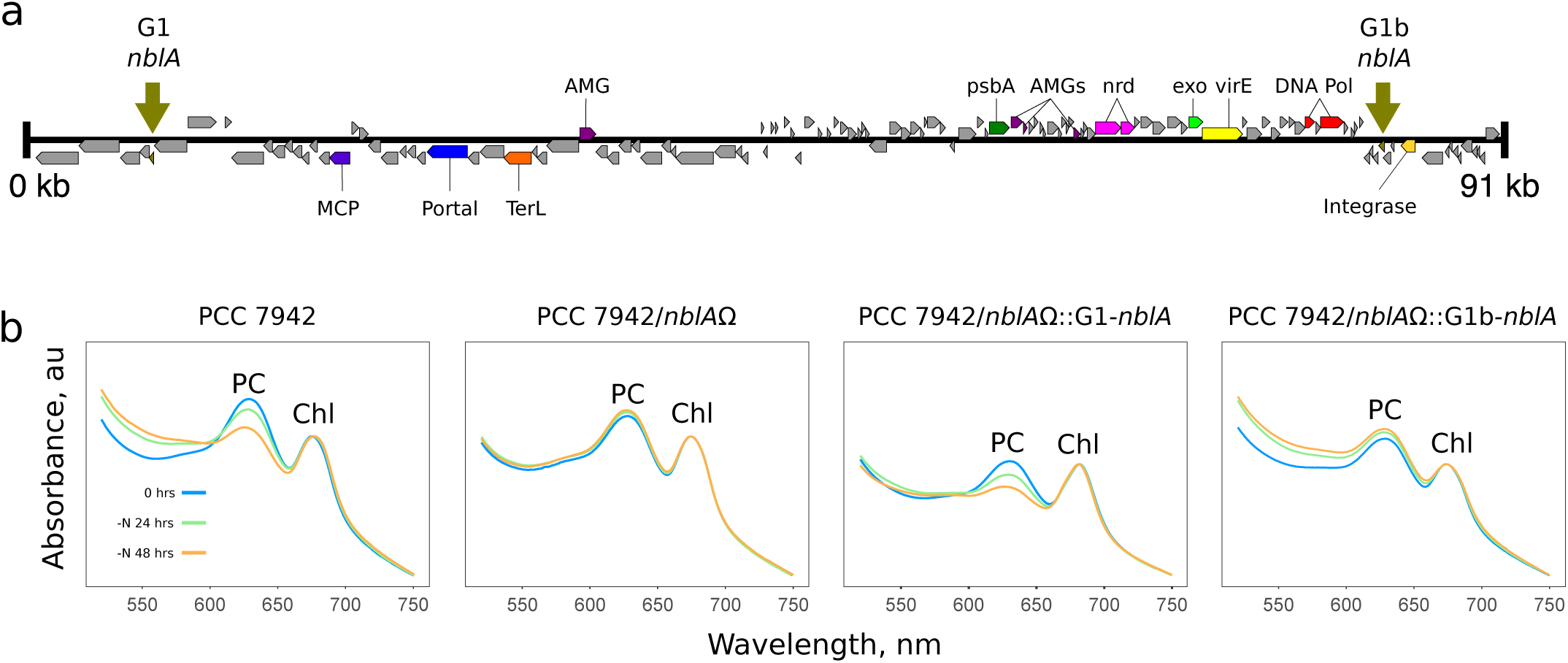
An uncultured marine cyanophage with two different *nblA*-like genes. **a.** Genome map of cyanophage IUI_154509. **b.** Absorbance spectra of *S. elongatus* PCC 7942 *nblA* knockout (PCC 7942/*nblA*Ω), Wild Type PCC 7942, PCC 7942 knockout complemented with phage G1 NblA (PCC 7942/*nblA*Ω::G1-*nblA*), and PCC 7942 knockout complemented with phage G1b NblA (PCC 7942/*nblA*Ω::G1b-*nblA*). Absorption spectra were recorded at the onset of nitrogen starvation time (0 h, blue line), and following 24 h (green line) and 48 h (orange line) nitrogen starvation (n=5). PC and chlorophyll-a (Chl) absorbance peaks are indicated.

In order to check if the NblA-like proteins encoded by cyanophage MAGs were indeed *bona fide* NblAs, we took advantage of a *S. elongatus* PCC 7942 NblA knockout strain (Sendersky et al. 2015; Collier and Grossman 1994). This mutant does not degrade its PBSs under nitrogen starvation and hence does not show the bleaching phenotype. We attempted to complement this mutant using the cyanophage NblA-like proteins under the control of the native *S. elongatus nblA* promoter. We chose two *nblA* genes (G1 and G1b) from a single cyanophage MAG (Figure 1a) for the complementation experiment. While in the G1-NblA variant degradation of PC was observed upon nitrogen starvation, the variant G1b-NblA was unable to restore the phenotype (Figure 1b).

To test whether the viral NblAs can potentially form dimers as cyanobacterial NblAs (Dines et al. 2008), we predicted their monomer and dimer structures using pyDockWEB (see Experimental Procedures for details). The structures of G1-NblA and G1b-NblA monomers were homology-modelled and their ability to form dimers was examined using pyDockWEB’s default parameters. G1-NblA demonstrated a homodimeric structure similar to cyanobacterial NblAs when the two monomers were docked (Figure S1). In contrast, the nonfunctional G1b-NblA did not form a homodimer, nor did the G1/G1b heterodimer. As a control we performed the same analysis for the reference cyanobacterial NblAs (PDB accessions 3CS5, 2Q8V and 1OJH), as well as for the two NblAs from *Synechocystis* sp. PCC 6803 that are known to form a heterodimer but were not crystallized (Baier et al. 2001; Baier et al. 2014). All of the known cyanobacterial dimers could be replicated using this *de novo* approach as formed the expected heterodimer for the two NblAs from *Synechocystis* (see Figure S1). These observations suggest that while G1-NblA is able to function as a homodimer as confirmed *in-silico*, G1b-NblA might require a different NblA to form a heterodimer, which can explain its inability to cause bleaching. One can tentatively hypothesize that this second monomer is not G1-NblA but an NblA provided by the yet unknown host of the virus.

In an initial analysis the putative NblA proteins from the MAGs appeared rather divergent from other proteins currently assigned in public databases to the NblA family (see Fig. 2c and Suppl. Fig. S2), therefore we collected a broader sample of NblAs and NblA-like proteins to understand their diversity. Combining protein profile with blast analysis and applying relaxed E-value thresholds to obtain divergent protein sequences in a taxonomy-independent manner, we collected protein sequences that were either annotated explicitly as NblA, NblA-like, Ycf18 or as proteins of unknown function from freshwater and marine cyanobacteria, red algae, and most intriguingly marine cyanophages alongside metagenomic assemblies (Fig. 2, Suppl. File 1 and 2 and Fig. S4). Eleven marine cyanophages, mostly cultured, appeared to contain single copies of *nlbA*, none of which is currently annotated as such (see Suppl. File 1 and Fig. S4). Moreover, we discovered four of these genes among unannotated ORFs. The additional metagenomic assembled contigs are of special interest, since they are annotated in GenBank as various unrelated bacterial (non-cyanobacterial) groups. A closer examination of the gene content and blast matches for the corresponding sequences revealed that all of them are in fact fragments of cyanophage genomes, except for one of likely cyanobacterial origin.

**Figure 2.**
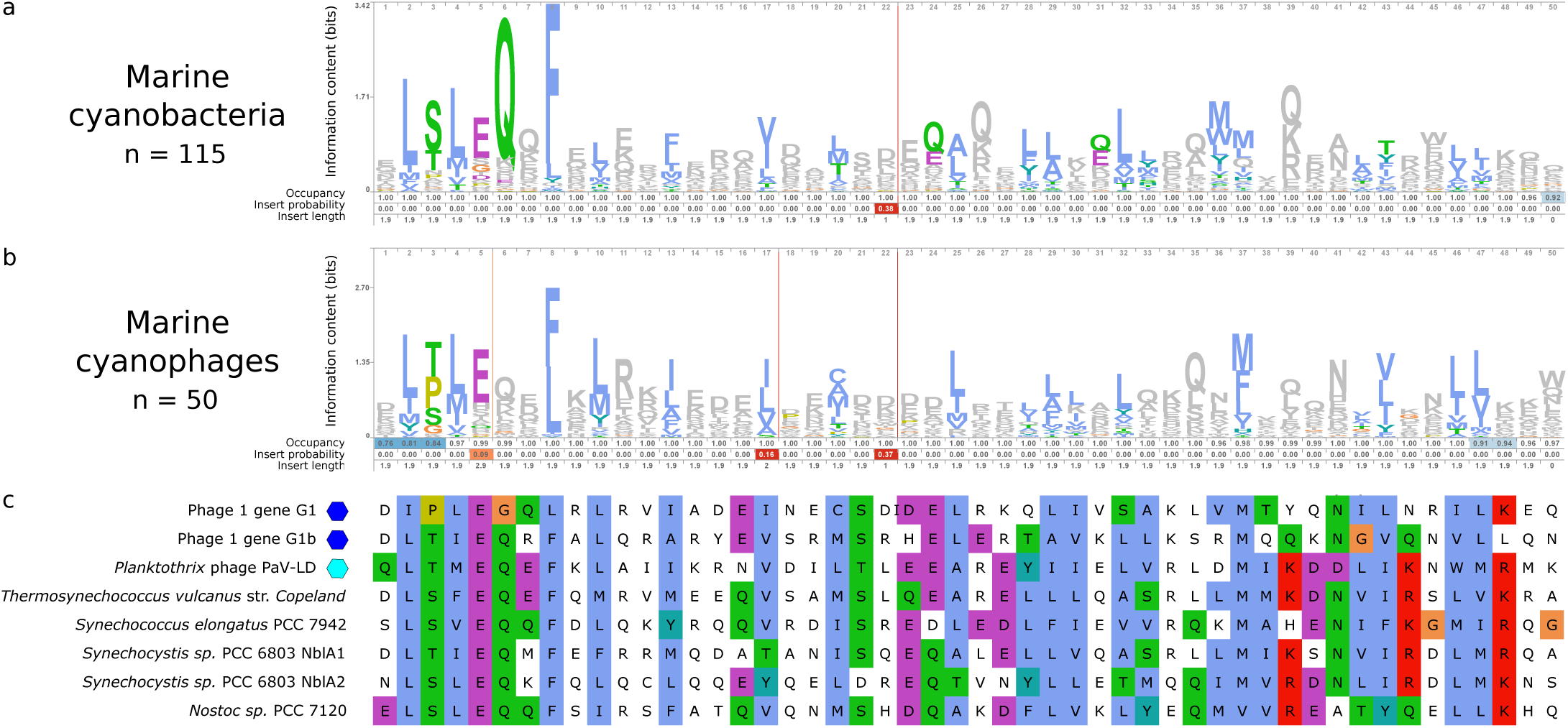
Protein profiles for the conserved region of NblA proteins. **a.** Protein profile for 115 NblA proteins from marine cyanobacteria. **b.** Protein profile for 50 NblA proteins derived from marine cyanophages (including metagenomic sequences erroneously annotated as various microbes). **c.** Fragment of the whole alignment of 871 NblA and NblA-like proteins showing chosen representatives: two marine cyanophage proteins investigated here (marked with blue diamonds) and NblAs studied in detail before, including a freshwater cyanophage (cyan diamond). All three panels are aligned to each other and trimmed to the boundaries of the Pfam NblA profile (PF04485). Amino acid properties and conservation relative to the corresponding profiles or whole alignment are highlighted according to the ClustalX color scheme.

Comparison of protein amino-acid profiles between marine phages and marine cyanobacteria revealed a higher overall diversity of the cyanophage NblAs, with most of the positions conserved in cyanobacteria conserved to a similar degree in their potential viruses (see Fig. 2a,b). Yet, some positions moderately or highly conserved in marine cyanobacteria (most prominently, Q6 in Fig. 2, cf. also Fig. 1 in (Dines et al. 2008)) allow more variability in marine cyanophages. These discrepancies between the two profiles explain the low scoring of marine cyanophage NblAs, including G1 and G1b, when matched against the NblA protein profile from Pfam (see Fig. S2) and their dissimilarity to the previously studied NblAs (see Fig. 2c).

Interestingly, unlike cyanophages, multiplicity of *nblA* genes seems to be the rule for cyanobacteria. Analysis of phycobilisome-containing cyanobacteria with chromosome-level genome assemblies indicated that most of them contain multiple *nblAs* (two and up to seven), with some *nblA* genes not being predicted or annotated, apparently due to their short sequence and high divergence (Suppl. Fig. S3).

Notwithstanding the short protein sequences and the corresponding alignment, we reconstructed a moderately supported phylogenetic tree for the protein family (see Suppl. Fig. S4). It should be stressed that due to the short alignment length, high sequence divergence and selection, the phylogenetic reconstruction is expected to be biased and is used here merely as a guide. Phage NblAs group into eight or nine putative clades, some of which are nested within cyanobacterial sequences indicating recent gene transfer from the corresponding hosts (e.g. *Phormidium* and *Planktothrix* phages) while others appear to branch deeply. The pairs of *nblA* genes from our cyanophage MAGs systematically belong to different divergent clades without apparent affinity to cyanobacterial NblAs (Figure 2). All of the NblAs from the apparently mis-classified phage contigs from metagenomic assemblies cluster with cyanophage NblAs. In line with the earlier observations by (Nakamura et al. 2013), the NblA-like proteins from red algae form two independent clades corresponding to the well-sampled plastid Ycf18 proteins and the smaller group of nuclear-encoded proteins (see Suppl. Fig. S4).

Our results show that, similarly to their freshwater relatives, *nblA* genes are carried by marine cyanophages. Moreover, one of the variants was successfully expressed in the heterologous *S. elongatus* model-system and was shown to be active. This implies that cyanophage NblAs play a role in the degradation of their hosts’ PC. As *S. elongatus* does not contain PE, future expression experiments with the viral G1 and G1b *nblA* gene variants in marine *Synechococcus* hosts containing both PC and PE are currently sought.

## Experimental Procedures

### Strains and culture conditions

*Synechococcus elongatus* PCC 7942 and derived strains were grown as described previously (Schatz et al. 2013). The cells were grown in the presence of antibiotics, and starvation was induced as described in Sendersky et al. (Sendersky et al. 2014).

### Marine cyanophage NblA detection

NblAs in the metagenomic assembled genomes were found with Hidden Markov Models trained on NblA amino acid sequences of freshwater cyanobacteria using HMMER and further analyzed employing HHblits in a pipeline based on HHpred (Remmert et al. 2011).

### Absorbance measurements

Absorbance was measured as described previously (Sendersky et al. 2014), with the exception that no integrating sphere was employed. Absorbance spectra were normalized at 680 nm.

### Cloning of *nblA*-like genes into pES57

Synthesized *nblA*-like genes (Hylabs, Rehovot, Israel), G1 and G1b were digested with *BamHI* and *NcoI*, cloned into pES57 and transformed into *Escherichia coli* DH10B. PCR with 5’-TGCAAGGCGATTAAGTTGG-3’ (forward) and 5’-TTGTGTGGAATTGTGAGCG-3’ (reverse) primers was used to verify the desired clones’ sequences.

### Structural predictions

Structural homology models for NblA proteins were built with SWISS-MODEL (Bienert et al. 2017). The pyDockWEB server (Jiménez-García et al. 2013) was used for *in silico* docking of the obtained NblA monomer structures. Three available crystal structures of cyanobacterial NblAs (PDB accessions 3CS5, 2Q8V and 1OJH) were recruited for reference. All structures were visualized in PyMOL (https://pymol.org/).

### Alignment and phylogenetic analysis

Sequences for phylogenetic analysis were collected by searching the Uniref90 database (https://www.uniprot.org/uniref/) with the Pfam NblA profile PF04485 (http://pfam.xfam.org/family/PF04485) using hmmsearch v. 3.1b2 (http://hmmer.org). According to empirical criteria, protein sequences with hits covering no less than 84% of the profile’s length were sampled with the default relaxed e-value threshold. The matching Uniref90 sequences, as well as remaining sequences assigned to the InterPro family IPR007574 and all Uniprot proteins with 50% identity to them were used as queries in a blastp v. 2.7.1+ (Altschul et al. 1990) search against Uniref90 with an e-value threshold of 0.03. The final dataset included all Uniprot KB sequences assigned to the matching Uniref90 and corresponding Uniref50 clusters, as well as the NblAs from the cyanophage MAGs analyzed here. In addition, a focused search for more divergent and/or unannotated *nblA*-genes was performed for all ORFs longer than 150 bp from cyanophage genomes available from Uniprot proteomes using the same strategy but with a blastp e-value threshold of 0.0001. All protein sequences (872 in total) were aligned with mafft v. 7.313 (Katoh and Standley 2013) in L-INS-i mode, trimmed with TrimAl v. 1.4.rev15 (Capella-Gutiérrez et al. 2009) (final alignment length of 55 positions) and the gene phylogeny was reconstructed with FastTree v. 2.1.9 (Price et al. 2010) under the LG-G model. One sequence (A0A2K8SZX4) had to be excluded due to misalignment at one of the flanks. Protein profiles for marine cyanobacteria and marine cyanophages were visualized with skylign (Wheeler et al. 2014).

### Gene number analysis

*nblA* gene numbers were inferred from cyanobacterial genomes tagged as complete in Genbank using the same strategy as described for the phylogenetic analysis above. As input, all predicted proteins and all ORFs at least 100 bp were taken and analyzed together.

## Supporting information

## Acknowledgments

We thank Faris Salama for help with the spectroscopic measurements. This work was funded by a European Commission ERC Advanced Grant (no. 321647), Israel Science Foundation grant (143/18) and the Louis and Lyra Richmond Memorial Chair in Life Sciences (to O.B.).

## Author Contributions

R.S. devised the initial idea for the project. O.N., S.L., R.S. and O.B conceived the experiments, J.F.-U. and A.R. performed bioinformatic analyses, and O.N. conducted the molecular biology and biophysical experiments. O.B. wrote the manuscript with contributions from all authors to data analysis, figure generation, and the final manuscript.

## Figures

**Supplementary Figure S1.**
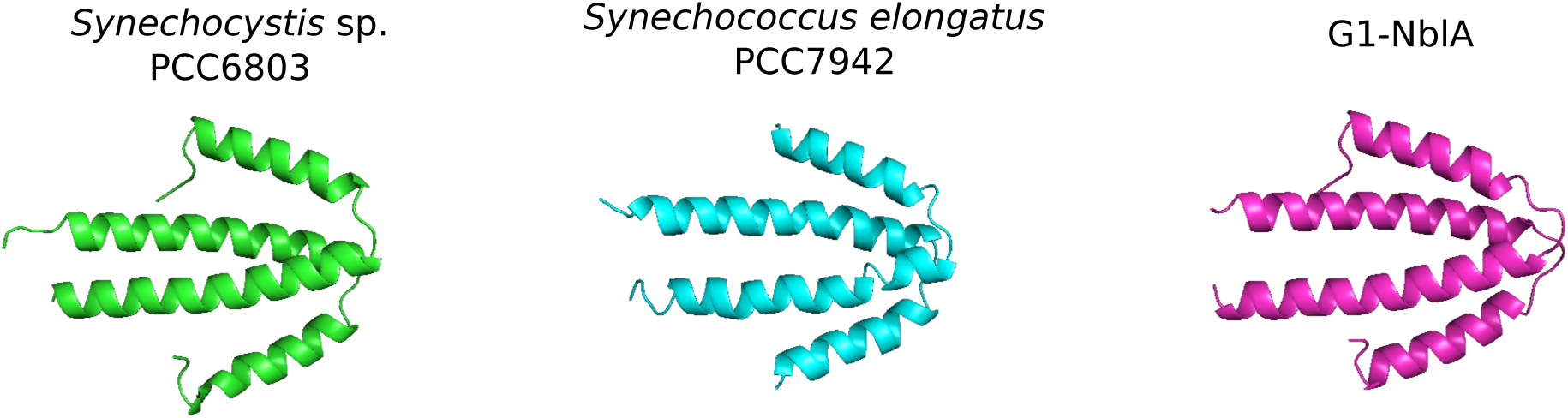
Structural predictions for NblA dimers from *Synechocystis* sp. PCC 6803 (P73890 and P73891 heterodimer), *Synechococcus elongatus* PCC 7942 (P35087 homodimer) and G1-NblA (homodimer) obtained with pyDockWEB.

**Supplementary Figure S2.**
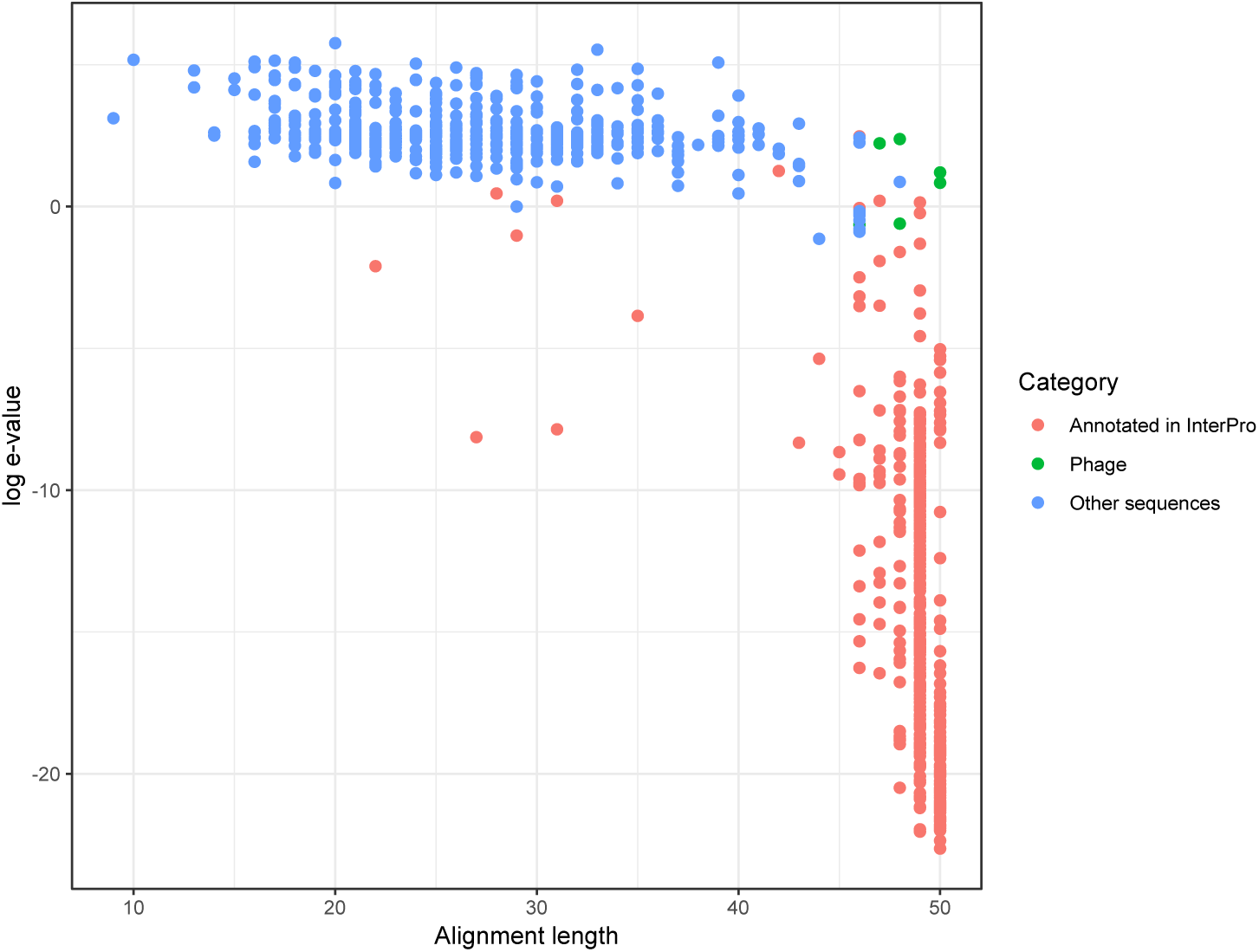
Distribution of e-values and alignment lengths of hmmsearch matches for the Uniref90 database and phage MAG when searched with the Pfam NblA profile PF04485. Uniref90 hits based on records outside of the Uniprot KB database are not shown.

**Supplementary Figure S3.**
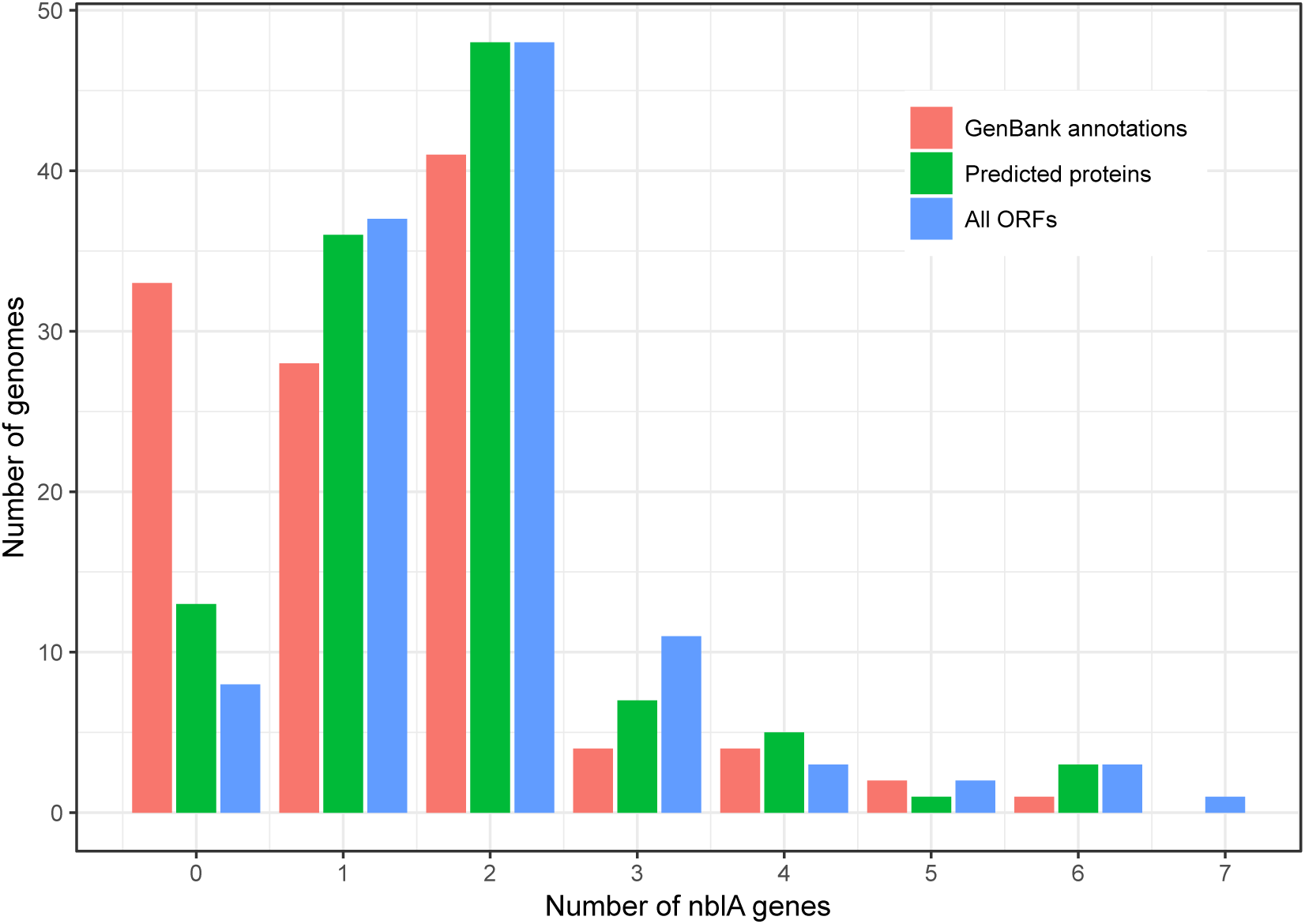
Frequency distribution of numbers of *nblA* genes in 126 complete genomes of cyanobacteria. Excluded are 13 *Prochlorococcus* genomes none of which encodes for phycobilisome proteins or *nblAs.* The counts provided correspond to three sets of annotations: explicit annotations from GenBank, predictions made by us for annotated proteins and predictions made for all ORFs of at least 100 bp long.

**Supplementary Figure S4.**
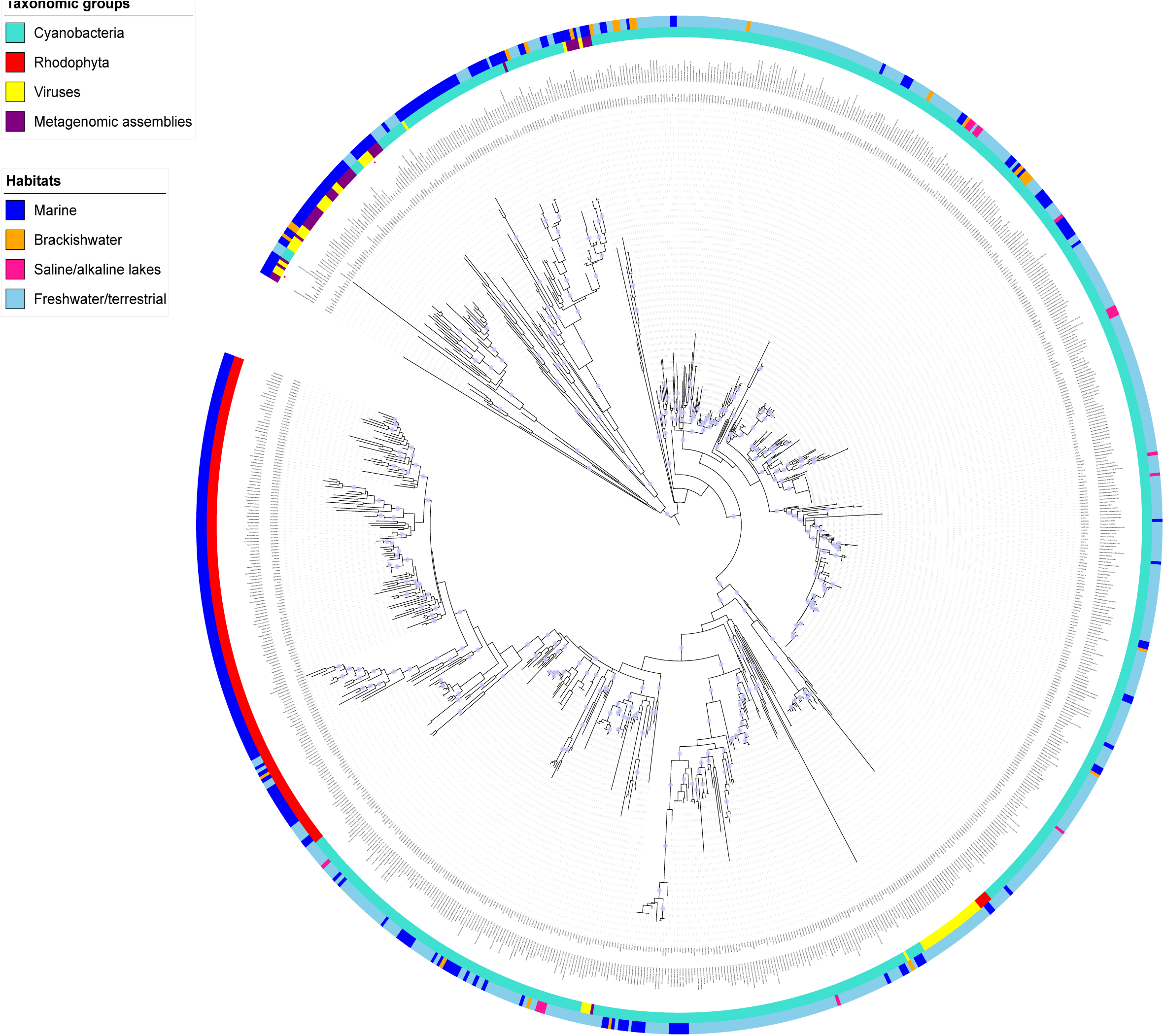
Phylogenetic tree of NblA, Ycf18 and NblA-like sequences. Branches with bootstrap support values of at least 80% are indicated with circles. G1-NblA and G1b-NblA are indicated with asterisks. Interactive representation of the tree is available at https://itol.embl.de/tree/13268108169281151544017427. The taxonomy assigned to the entries is left unchanged except in the following cases: all non-cyanobacterial metagenomic assemblies are grouped into one category; L8AI55 and L8AJF3 are derived from a *Synechocystis-Bacillus* chimera; A0A2R7TRJ3 is apparently from a short fragment of a *Synechococcus* genome mis-classified as *Pseudomonas* sp. HMWF031.

**Supplementary File 1** (spreadsheet). Metadata for the 872 NblA and NblA-like proteins collected by us for the phylogenetic analysis and protein profiles, including marine cyanophages analyzed here.

**Supplementary File 2** (fasta file). Untrimmed alignment of the NblA and NblA-like proteins.

